# Curvature increases permeability of the plasma membrane for ions, water and the anti-cancer drugs cisplatin and gemcitabine

**DOI:** 10.1101/602177

**Authors:** Semen Yesylevskyy, Timothée Rivel, Christophe Ramseyer

## Abstract

In this work the permeability of a model asymmetric plasma membrane, for ions, water and the anti-cancer drugs cisplatin and gemcitabine is studied by means of all-atom molecular dynamics simulations. It is shown that permeability of the membranes increases from one to three orders of magnitude upon membrane bending depending on the compound and the sign of curvature. Our results show that the membrane curvature is an important factor which should be considered during evaluation of drug translocation.

**TOC GRAPHICS:** 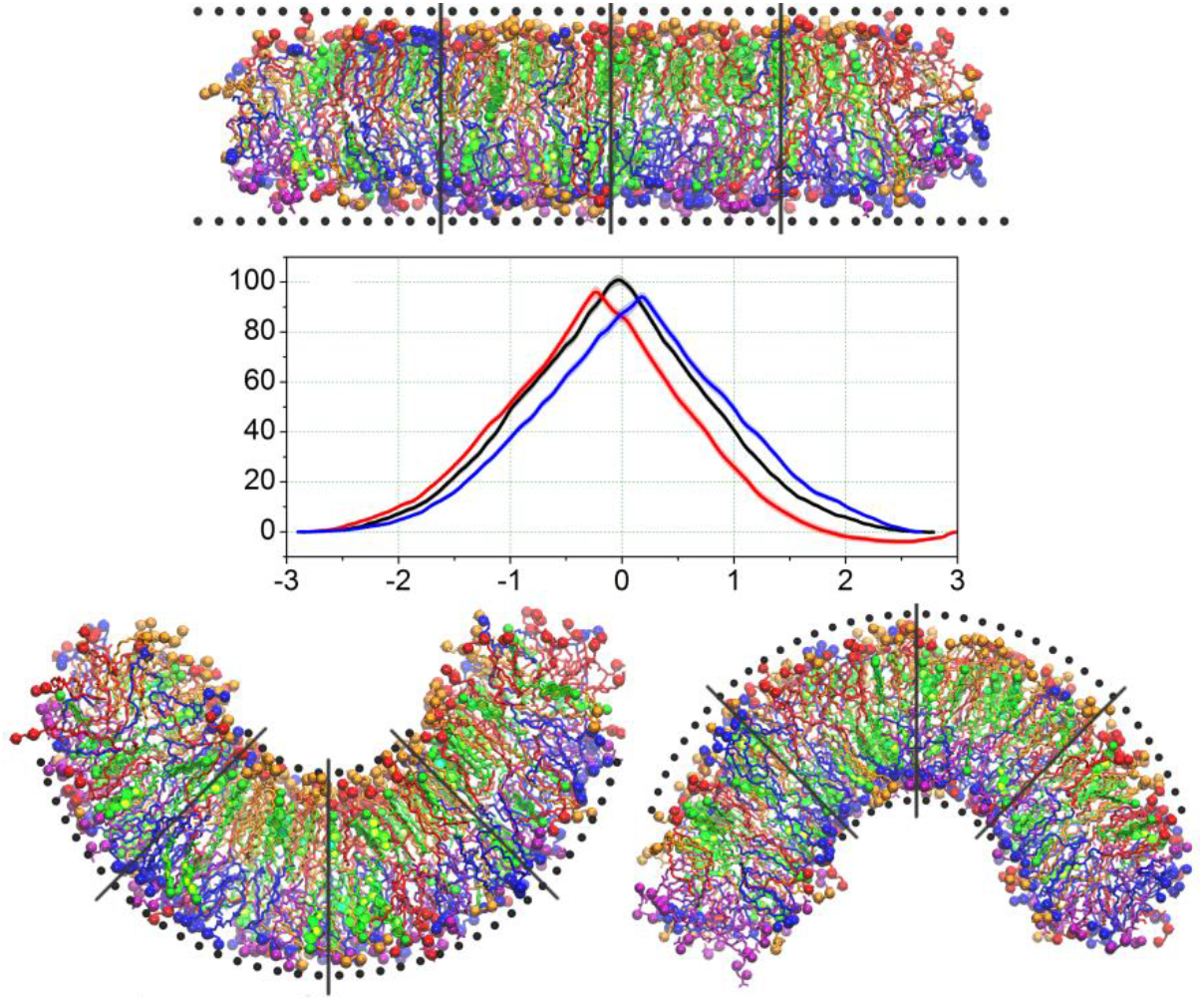

---

Passive permeability of cell membranes is of great importance for pharmacology since many drugs are known to permeate spontaneously through the plasma membrane of the target cells. The membranes of real cells are highly heterogeneous and possess regions with very different curvature – from essentially flat to highly curved, such as filopodia, cilia, membrane blebs, caveolae, etc^1^. Origins of the membrane curvature are diverse and range from spontaneous curvature of the lipids ^2–4^ to the influence of membrane proteins and cytoskeleton ^5^. Membrane curvature is intrinsically connected to the asymmetry of lipid ^6–7^ and cholesterol ^8–10^ content of membrane monolayers.

Up to now it is not known which regions of the membrane are preferred for drug permeation and whether such preference exists. Moreover, membrane morphologies of normal and malignant cells differ substantially ^11–12^. Malignant cells could have more ragged surface with many membrane blebs and protrusions or, in contrast, be round and featureless in comparison to their normal counterparts. It is not known whether these differences in membrane morphology affects uptake of the drugs by cancer cells. That is why in this work we consider two widely used drugs – cisplatin and gemcitabine.

Being a transient dynamic property of the membrane, which changes in time and in different sites of the cell surface, the curvature is difficult to study experimentally. Particularly studying the influence of curvature on the passive transport of small molecules and drugs through the membrane remains challenging.

Molecular dynamics (MD) simulations allow overcoming these difficulties by providing atomistic view of the membrane in completely controllable environment. Recent MD study revealed significant and non-trivial effects of curvature on the distribution of cholesterol in the membrane, on the thickness of its leaflets and on the order parameter of the lipid tails ^13–14^. MD simulations also provide unique opportunity of studying diffusion of drugs and small molecules through realistic model membranes in atomic details (see Supporting Information for discussion of available methodologies and their limitations).

In this work we used all-atom MD simulations for determining permeability of curved membranes for water, ions and small hydrophilic anti-cancer drugs cisplatin and gemcitabine. We used a model of mammalian plasma membrane with asymmetric lipid composition and 33% mole fraction of cholesterol. We performed simulations for three values of curvature: 0 nm^−1^ (flat membrane), 0.2 nm^−1^ (outer monolayer is convex) and −0.2 nm^−1^ (inner monolayer is convex) as shown in Fig.1. The potentials of mean force (PMFs), diffusion coefficients (*D*) and permeation resistances (*R*) across the membrane were computed. The dependence of permeability (*P*) on the curvature for all studied ligands is estimated.

**Figure 1.**
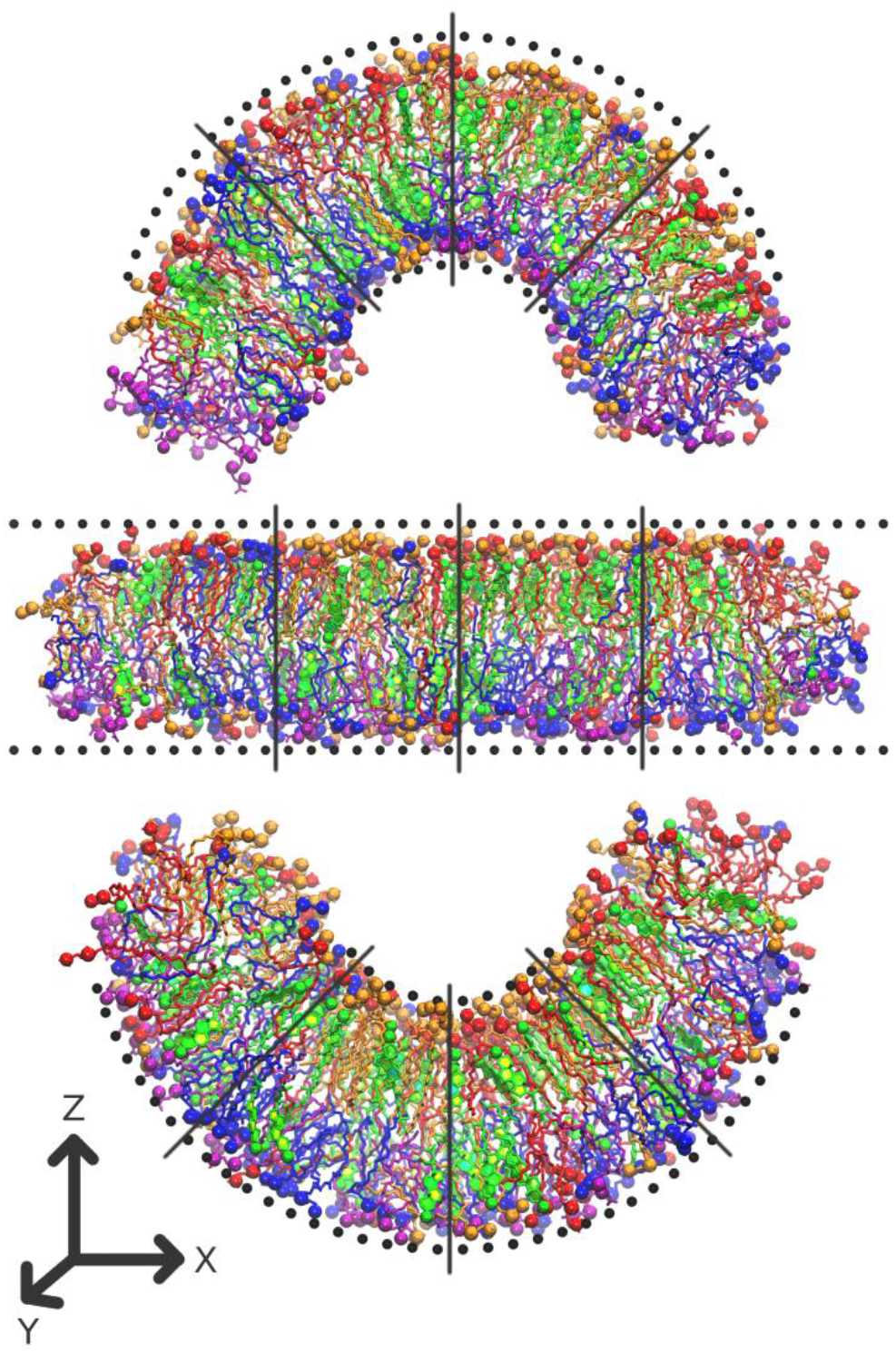
Simulated systems with the curvatures 0.2 nm^−1^ (top panel), 0.0 nm^−1^ (middle panel) and −0.2 nm^−1^ (bottom panel). Outer membrane leaflet is on top. PC is red, PE is blue, PS is violet, SM is orange and cholesterol is green (see Methods for definitions of abbreviations). N and P atoms of the lipid head group and the hydroxyl oxygen of cholesterol are shown as spheres. Black spheres show dummy particles which maintain the membrane shape. Black lines show approximate axes where the ligands are restrained during umbrella sampling simulations. Water molecules and ions are not shown for clarity.

Analysis of the PMFs shows that the energy barriers for all permeating ligands are lower in the curved membrane in comparison to the flat one (Fig. 2) while the magnitude of this effect varies between the ligands (Table 1).

**Figure 2.**
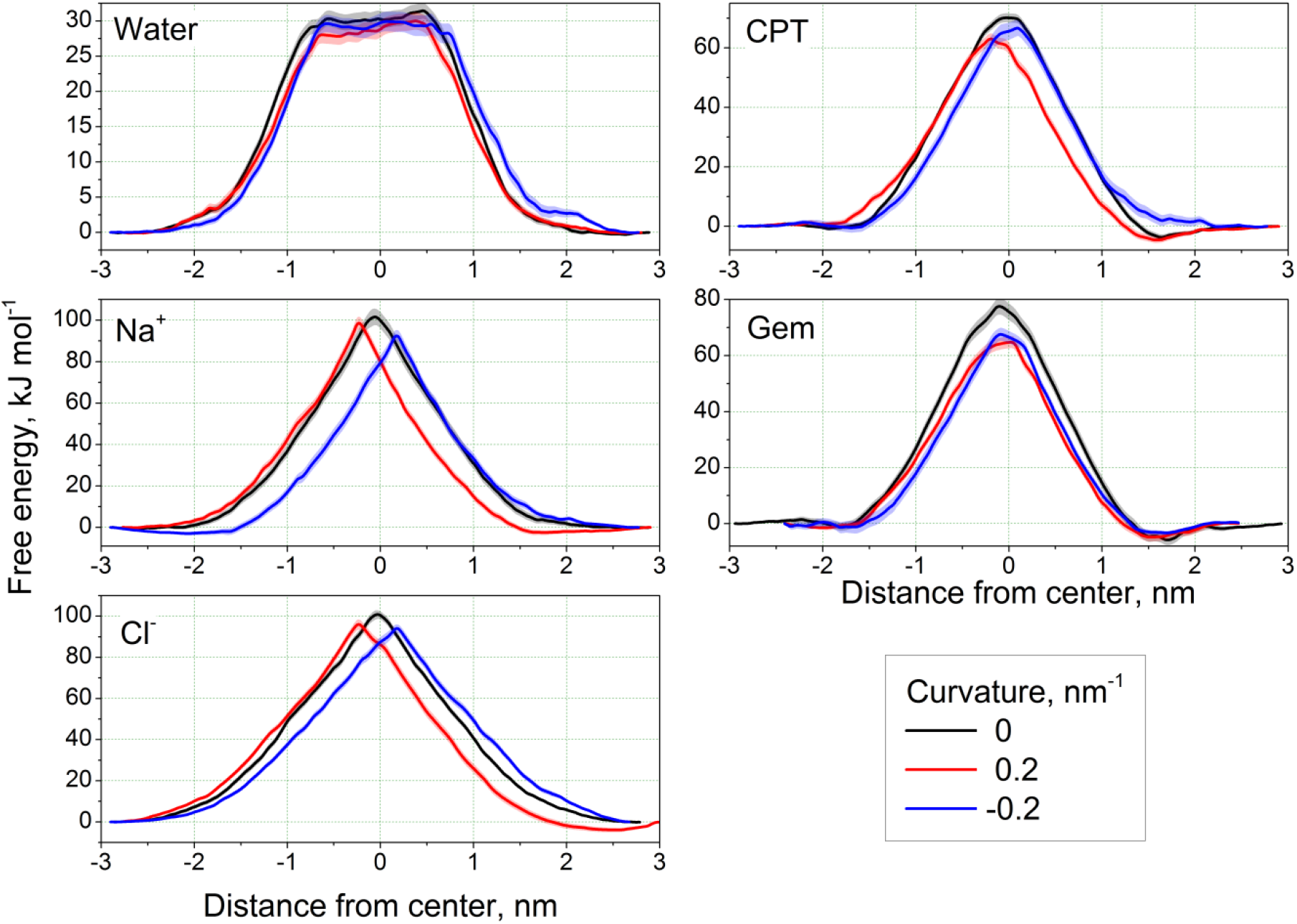
Potentials of mean force for studied ligands. The error bars are shown as semi-transparent bands.

**Table 1.** Maximal heights of the energy barriers (kJ/mol). Values in parentheses show relative height of the barrier with respect to the flat membrane for particular ligand.

| Curvature | Na<sup>+</sup> | Cl<sup>-</sup> | Water | CPT | Gem |
| --- | --- | --- | --- | --- | --- |
| 0 | 101.1 | 101.6 | 31.4 | 70.1 | 77.5 |
| 0.2 | 98.3 (0.97) | 96.4 (0.95) | 30.2 (0.96) | 63.1 (0.90) | 64.8 (0.83) |
| -0.2 | 92.7 (0.92) | 93.4 (0.92) | 29.9 (0.95) | 66.7 (0.95) | 67.6 (0.87) |

The peaks of the PMFs shift systematically towards concave monolayer (Figure S1, see Supporting Information for definition of the peak position). Such shift is expected since the convex monolayer has larger area per lipid and looser packing of the lipids than the flat membrane while the concave monolayer is more densely packed ^13^ and imposes higher barrier on the permeant.

The diffusion coefficients of the ligands change consistently upon the membrane bending (Fig. 3). For all ligands except Gem the *D* is lower for the concave monolayer in comparison to the flat membrane which reflects more congested environment in this monolayer. In the case of Gem the changes of *D* are small and do not allow to deduce obvious trend from them.

**Figure 3.**
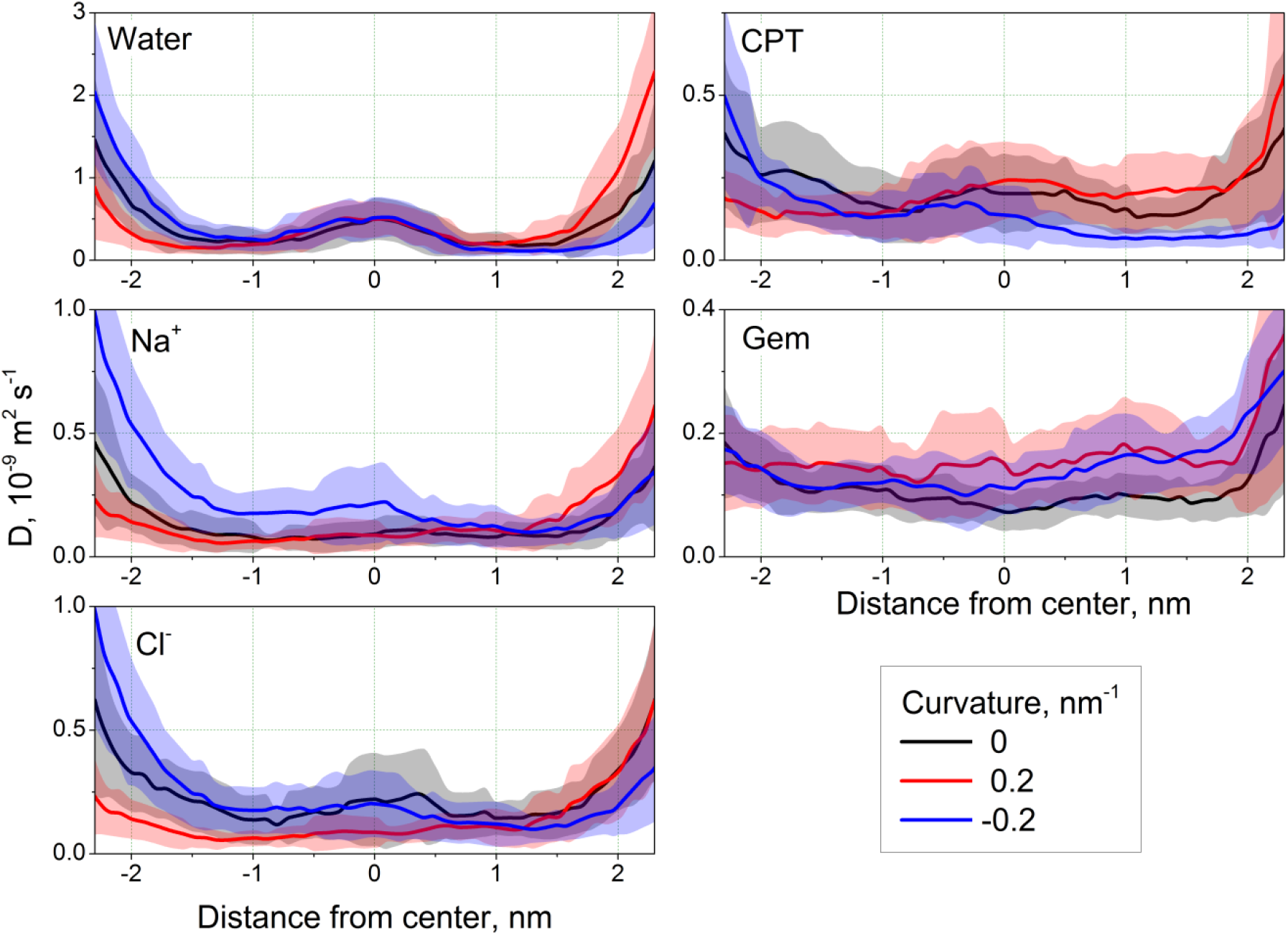
Diffusion coefficients of the studied ligands. The error bars are shown as semi-transparent bands.

The absolute values of *D* do not change much upon bending within 1 nm from the center of the membrane. The maximal change in this area is about 1.5 times for all ligands. Much larger variations are observed in the regions of lipid head groups and bulk solvent. In order to estimate the influence of these regions on total permeability we analyzed the membrane resistance *R* (Figure S2). For all studied ligands resistance drops by several orders of magnitude at the distance of ~1 nm from the membrane center (note the log scale in Fig. S2) thus the changes of *D* outside this region has almost no effect on permeability. This suggests that the permeability will depend only slightly on *D* and the shape and height of the PMF plays the major role.

Permeability coefficients *P* of the ligands are summarized in Table 2. There is a systematic trend, which is consistent with the changes of PMFs upon bending. The *P* of all ligands increases upon membrane bending. For Na^+^ and Cl^−^ ions this increase is much larger for negative curvature (when inner monolayer in convex). For water the effect is comparable for positive and negative curvatures. For CPT and Gem the decrease of *P* is much larger for positive curvature (when inner monolayer in concave). Comparison with Table1 shows that the permeability follows exactly the same trend as the decrease of the energy barrier observed on PMFs in curved membranes.

**Table 2.** Permeability coefficients (m·s^−1^) for all studied ligands. Numbers in parentheses show the ratio of permeability in comparison to zero curvature for particular ligand.

| Curvature, $\text{nm}^{-1}$ | 0 | 0.2 | -0.2 |
| --- | --- | --- | --- |
| Water | $2.5\cdot 10^{-6} \pm 3.6\cdot 10^{-8}$ | $5.5\cdot 10^{-6} \pm 7.2\cdot 10^{-8}$<br>(2.1) | $3.7\cdot 10^{-6} \pm 5.3\cdot 10^{-8}$<br>(1.5) |
| $\text{Na}^+$ | $1.5\cdot 10^{-17} \pm 1.2\cdot 10^{-18}$ | $8.4\cdot 10^{-17} \pm 6.9\cdot 10^{-18}$<br>(5.3) | $1.7\cdot 10^{-15} \pm 1.3\cdot 10^{-16}$<br>(107.5) |
| $\text{Cl}^-$ | $4.6\cdot 10^{-17} \pm 2.4\cdot 10^{-18}$ | $1.5\cdot 10^{-16} \pm 8.7\cdot 10^{-18}$<br>(3.2) | $6.4\cdot 10^{-16} \pm 3.3\cdot 10^{-17}$<br>(13.8) |
| CPT | $2.3\cdot 10^{-12} \pm 6.2\cdot 10^{-14}$ | $5.0\cdot 10^{-11} \pm 1.6\cdot 10^{-12}$<br>(22.1) | $7.0\cdot 10^{-12} \pm 3.1\cdot 10^{-13}$<br>(3.0) |
| Gem | $8.3 \cdot 10^{-14} \pm 3.6 \cdot 10^{-15}$ | $1.4 \cdot 10^{-11} \pm 4.2 \cdot 10^{-13}$<br>(164.5) | $4.5 \cdot 10^{-12} \pm 1.5 \cdot 10^{-13}$<br>(53.7) |

There is no obvious dependence of the absolute change of permeability on the size or chemical nature of the ligand. For example, the absolute change for Na^+^ ions is one order of magnitude larger than for Cl^−^ ions for positive curvature, but they are of the same order of magnitude for the negative curvature.

The main message which could be inferred from our data is that the curvature affects permeability of the realistic asymmetric membranes and decreases it up to three orders of magnitude depending on the direction of bending and the permeating ligand. Particularly, the difference in permeability in respect to the flat membrane may reach ~160 times for gemcitabine and ~22 times for cisplatin. This allows us speculating that in real cancer cells most of the passive transport of these drugs is likely to occur in highly curved regions of the membrane. Thus the changes of membrane curvature in malignant cells may contribute to drug resistance mechanisms.

## Conclusions

It is shown that permeability of the asymmetric lipid membrane, which mimics the composition of mammalian plasma membrane, for ions, water, cisplatin and gemcitabine strongly depends on the curvature. The permeabilities of all studied compounds increase from one to three orders of magnitude upon membrane bending depending on the permeating compound and the sign of curvature. Our results show that the membrane curvature could not be neglected during evaluation of the membrane permeability for hydrophilic drugs. Modulation of the membrane curvature in cancer cells may play a role in the mechanisms of drug resistance due to curvature-dependent permeability of the common anti-cancer drugs.

## Computational Methods

All simulations were performed using GROMACS software ^15^, versions 5.1.2 and 2018.2. The slipids force field ^16^ is used for lipids in combination with AMBER99sb force field for water and the ions. All simulations were performed in NPT conditions with Berendsen barostat ^17^ at 1 atm and velocity rescale thermostat ^18^ at 320 K. The group cutoff scheme was used ^19^. An integration step of 1 fs was used for cisplatin (as required by its flexible topology ^20^) and 2 fs for all other ligands. For simulations with 2 fs time step all bonds were converted to rigid constraints. Long range electrostatics was computed with the PME method ^21^. Preparation of the systems and data analysis was performed with Pteros 2.0 molecular modeling library ^22–23^. VMD 1.9.2 was used for visualization ^24^.

Topologies of water and ions were taken from AMBER99sb force field. Topologies of cisplatin and gemcitabine were developed and tested in our previous works ^20, 25^ respectively. The QDSol topology of cisplatin was used ^20^.

The simulation setup for curved asymmetric membranes developed in our previous works ^13, 26^ was used. The membrane is prepared as a bicelle which is limited by semi-cylindrical caps in XZ plane and forms an infinite bilayer in Y direction (Fig. 1). When the curvature is imposed on the membrane the areas of concave and convex monolayers become different and the lipids redistribute through the caps of the bicelle in order to compensate for resulting mechanical strain. The caps serve as “compensating reservoirs” which store excessive lipids from concave monolayer and donate the lipids to convex monolayer. This setup has shown robust performance in both coarse-grained and atomistic simulations ^13, 26^.

Table 3 shows the lipid content of the monolayers designed according to well-established lipid content of mammalian erythrocyte membranes ^27^. Phosphatidylinositol was not included into simulations due to its small concentration, which results in about one molecule per system.

**Table 3.** Lipid content (absolute number of molecules) of the monolayers. Initial cholesterol distribution is shown.

| Component | Outer monolayer | Inner monolayer |
| --- | --- | --- |
| SM (sphingomyelin) | 42 | 12 |
| PC (1,2-dioleoyl-sn-glycero-3-phosphocholine) | 46 | 14 |
| PE (1,2-dioleoyl-sn-glycero-3-phosphoethanolamine) | 14 | 46 |
| PS (1,2-dioleoyl-sn-glycero-3-phospho-L-serine) | 0 | 30 |
| Cholesterol | 51 | 51 |

The mixing of the lipids from different monolayers in the regions of bicelle caps is prevented by imposing selective artificial repulsive potentials developed in our previous works ^13, 26, 28–29^.

We use the method of keeping global membrane curvature at given value by shaping the membrane by means of dummy particles ^13^. In brief the idea is to put the membrane between two repulsive surfaces of artificial particles (the walls) which scaffold the global shape of membrane but do not affect lateral dynamics of individual lipids.

The system is pre-equilibrated as a bilayer with periodic boundary conditions for 10 ns and then converted to a bicelle by adding extra layers of water from both sides in X direction. After that the restricting walls are added and the system is equilibrated as a planar bicelle for ~250 ns.

The inhomogeneous diffusion (ISD) model was used to compute permeabilities of the ligands. The umbrella sampling simulations were used to compute the potentials of means force and the diffusion coefficients of the ligands across the membrane.

The details for all the steps of system preparation, simulation, analysis and error estimation are provided in Supplementary Information.

## ASSOCIATED CONTENT

### Supporting Information

The following files are available free of charge.

Detailed computational methods and additional results (PDF).

## AUTHOR INFORMATION

CR and SY designed the study. SY developed simulation methodology. SY and TR developed software for data analysis, performed simulations, analyzed and visualized data. CR guided discussion and interpretation of the results. The manuscript was written by SY and CR. The authors declare no competing financial interests.

## ACKNOWLEDGMENT

This work was supported by the European Union’s Horizon 2020 research and innovation programme under the Marie Skłodowska-Curie grant agreement No. 690853. The computational studies work was performed using HPC resources from GENCI- [TGCC/CINES/IDRIS] (Grant 2018-[A0050710578]), the Centre de calcul Champagne-Ardenne Romeo and the Mésocentre de calcul de Franche-Comté.

