## Supplementary information for "Curvature increases permeability of the plasma membrane for ions, water and the anti-cancer drugs cisplatin and gemcitabine"

#### ADDITIONAL INFORMATION FOR INTRODUCTION

*Methodology of computing permeability and its limitations*

The methodology of computing permeabilities from MD simulations is still subject to debate. There is a general consensus that obtaining precise quantitative values of permeabilities from MD simulations is an extremely complex task which requires further methodological development. Particularly, quantitative comparison of permeabilities of different ligands in MD simulations, which is very important for the drug design studies, is currently considered unreliable. However, MD simulations could still be used with great success to compare the permeabilities of the same ligand in different conditions on the semi-quantitative level. In this case identical systematic errors apply to all simulations, which make them directly comparable to each other despite the fact that exact values of permeabilities are unlikely to be sufficiently accurate.

The most widespread method of computing permeability is the inhomogeneous solubility-diffusion (ISD) model [1-2], which relies on the knowledge of non-uniform diffusion coefficient of the ligand across the membrane. Although the ISD model was used successfully in many studies [3-5] the systematic difficulties in computing free energy profiles and diffusion coefficients are reported [3, 6-10]. One of the concerns comes from the difficulty of determining equilibrium state of the ligand in the membrane related to very slow transitions in bilayer structure, which occur on the time scale beyond the capabilities of all-atom MD simulations [11]. Another concern is related to anomalous diffusion of the ligands on typical MD time scales [12]. The influence of the longlived correlations of the restrained ligand on diffusion coefficients in the membrane was also suggested as a possible source of systematic errors in ISD [13].

In our study we rely on pragmatic approach, which allows making conclusions on semi-quantitative level. We never compare different ligands with each other in terms of absolute values of permeability or translocation energy barriers. However, we compare the values for the same ligand in the same membrane but with different curvatures. Any systematic errors should be very similar in such simulations, which makes them directly comparable to each other.

We are aware that it is not possible to achieve true equilibrium sampling in complex multi-component membrane at the time scale of hundreds of nanoseconds. Simulation times of hundreds of microseconds or even milliseconds are needed for this, which is beyond the capabilities of modern computers for all-atom simulations. Thus, our simulations only sample some local metastable states of the membrane. We computed PMFs for several ligands in different parts of the membrane simultaneously which allows sampling several different microenvironments of the ligand. If reaction of these microenvironments to the membrane bending is significantly different, then averaging over multiple ligands in the same simulation will either lead to unstable and poorly converged PMF or the trends for different compounds would be inconsistent. However, we obtain well converged PMFs and consistent trends for all studied compounds. This means that the influence of curvature on all sampled microenvironments of the ligands is very similar and the physical effect of curvature is caught correctly in our simulations.

In contrast to the PMFs, the results for diffusion coefficients are less robust. Despite averaging over multiple ligands and additional smoothing, the curves for $D$ are still noisy. The influence of curvature is well detectable in general but not in fine details. However, permeability depends exponentially on PMF and linearly on $D$, thus even large uncertainties in the values of $D$ could be tolerated without changing qualitative results.

It is possible to conclude that better sampling will probably influence the results quantitatively but it is unlikely that the qualitative trends will change.

*Approaches to maintaining membrane curvature in simulations*

In order to study the influence of curvature on permeability one has to sample the transmembrane diffusion of the ligand in the parts of the membrane with desired curvature only. There are two ways of achieving this: either by sampling large membrane patch for a long time and selecting the regions with needed curvature, utilizing one of known methods of determining local [14] or global [15] membrane curvatures, or by maintaining the curvature of the membrane around desired value by artificial restrains. The first approach, while being closer to reality, is currently well beyond the time scales available for all-atom MD simulations. In contrast, the second approach is currently computationally tractable. In our previous work [16] we developed an efficient technique of restricting global membrane curvature to any desired value, which does not affect fine-grained dynamics of membrane components and key macroscopic membrane properties.

### ADDITIONAL RESULTS

*Peak positions on the PMFs*

The position of the peak on the PMF is computed as the first moments of the PMF:

$$m_1 = \frac{\int_{z_2}^{z_1} zW(z)dz}{\int_{z_2}^{z_1} W(z)dz},$$

where $z_1$ and $z_2$ are the limits in which the PMF W(z) is computed.

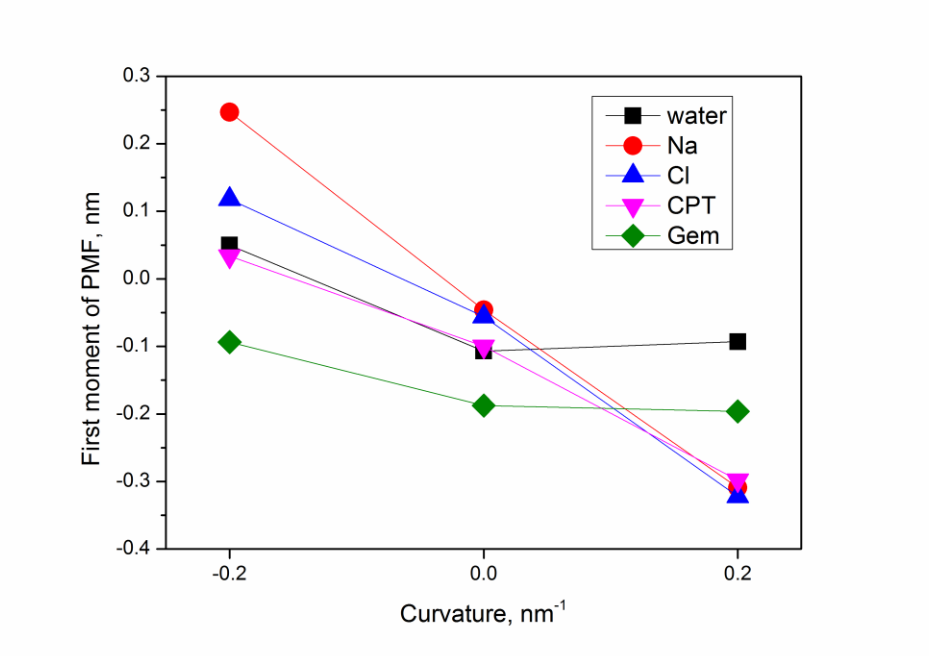


**Figure S1.** Positions of the peaks of PMFs, computed as the first moments of the corresponding curves, as a function of curvature.

*Membrane resistance profiles*

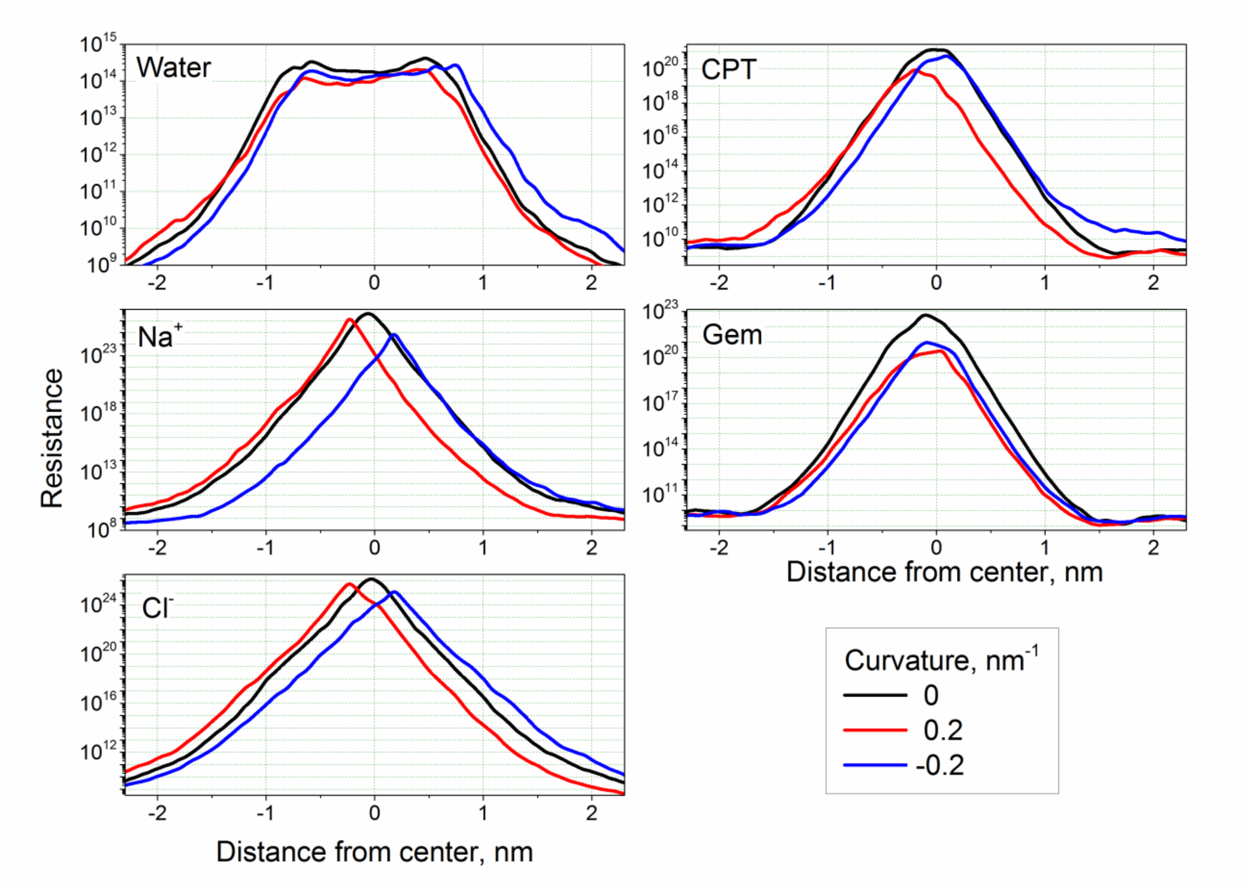

**Figure S2.** Permeation resistance R of PM membrane for all studied ligands for different curvature.

### DETAILED COMPUTATIONAL METHODS

*Preventing mixing of the monolayers*

All heavy atoms of the lipids (except the carbon atoms of the distal parts of the tails below the double bond) are assigned an additional Van der Waals repulsive potential (σ=0.8 nm, ε=1·10-7 J/mol) which acts between the lipids of inner and outer monolayers only. This allows the ends of the lipid tails to interact normally in the bilayer region of the bicelle while the lateral contacts between the lipids from different monolayers in the caps of the bicelle become energetically unfavorable. Such setup prevents the mixing of the monolayers effectively without any detectable effect on the bilayer part of the system. Cholesterol molecules diffuse freely through the caps of the bicelle which facilitates their optimal distribution between the monolayers.

*Membrane bending procedure*

The scaffolding walls only interact with carbon atoms of the lipid tails by means of Van der Waals interactions. Since the walls are impermeable for the hydrophobic lipid tails but completely transparent for all other atoms they restrict the general shape of the membrane effectively without influencing lateral diffusion of the lipids and dynamics of lipid head groups, cholesterol, ions, water molecules and ligands.

The planar walls are initially built around the bicelle with zero curvature. After that the shape of the walls is changed gradually by slowly pulling their beads by external forces towards new positions. The dummy particles push the hydrophobic core of the membrane and force it to bend accordingly. After the bending the membrane is equilibrated for 250 ns for each target curvature prior to production simulations.

The walls are composed from independent non-interacting beads which consist of anchor and shell particles. Anchor is a dummy particle which is fixed in absolute coordinates and does not interact with any other particles in the system. The shell particle is connected to an anchor by harmonic bond of zero length with the force constant of 10000 J·mol-1·nm-2. The shell particles interact selectively with the carbon atoms of the lipid acyl tails by means of Van der Waals interactions only. Parameters of this Lennard-Jones interaction are σ=0.85 nm, ε=1·10-5 J/mol which means that the potential is purely repulsive within the short-range cut-off rc=0.8 nm adopted in Amber force filed. Such setup eliminates unwanted direct interactions of the lipid atoms with fixed anchor particles and allows positioning the walls precisely at the same time.

The walls are positioned initially at the level of lipid head groups of each monolayer of the flat bilayer at the distance of 5 nm from each other. The dummy beads are arranged at rectangular grid in the plane of the wall with the spacing of ~0.51 nm.

Since the walls are impermeable for the hydrophobic lipid tails but completely transparent for all other atoms they restrict the general shape of the membrane effectively without influencing lateral diffusion of the lipids and dynamics of lipid head groups, ions and water molecules. The distance between the walls and the parameters of repulsive Van der Waals interactions are adjusted empirically in order to minimize the influence of the walls on membrane properties. The density profiles of various chemical groups of the lipids were compared for planar bilayers with and without the walls. Obtained results show that the changes of bilayer structure introduced by the walls are barely visible and could be neglected safely [16].

In order to produce the curved membranes the following procedure is used. The planar bilayer equilibrated at the presence of the walls is used as initial system. The desired curvature ρ is set and the center of curvature is determined as shown in Fig. S3. New positions of the anchor particles, which correspond to desired curvature, are computed. The distances between the dummy beads within each wall are kept constant to ensure that the density of dummy particles per unit area of each monolayer is the same as in the planar bilayer. After that all anchor particles are moved with the constant rate towards their new positions during 5 ns. The dummy particles push the hydrophobic core of the membrane and force it to bend accordingly. Due to different arch length of upper and lower walls some beads of the later, which appear beyond the needed sector, are removed as shown in Fig S3. After that the system is equilibrated for at least 200 ns with the anchor particles fixes at new positions to relax any tension introduced by the forced bending. The bending is performed in several stages with the curvature step of 0.05 $nm^{-1}$ (from 0 to 0.05 $nm^{-1}$, from 0.05 to 0.1 $nm^{-1}$, etc.) to reduce mechanical stress imposed on the bicelle on each stage. For our asymmetric bicelle the direction of bending (upwards or downwards) matters due to different lipid content of monolayers. The systems with final curvatures (0.2 $nm^{-1}$ and -0.2 $nm^{-1}$) were additionally equilibrated for 250 ns.

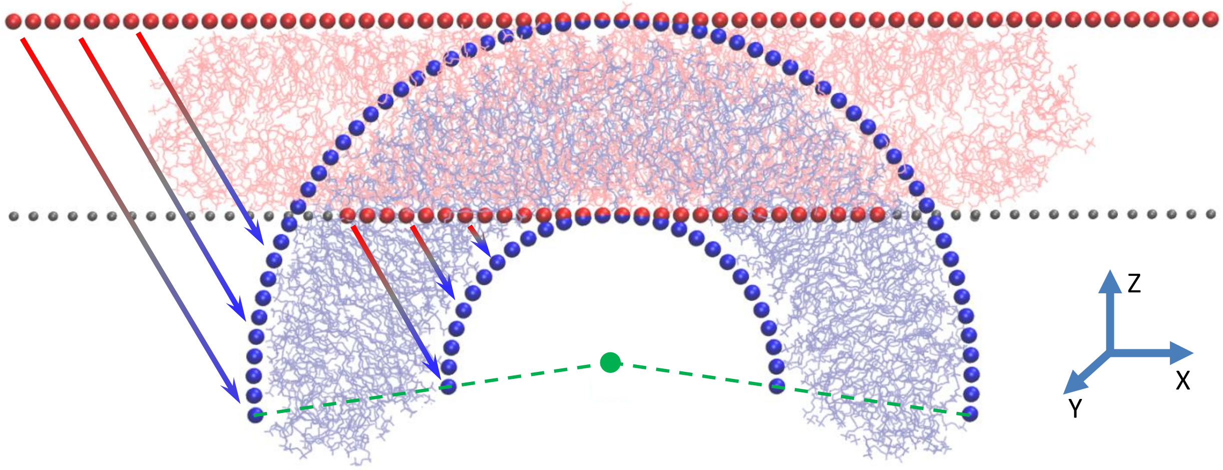


**Figure S3.**The scheme of membrane bending. The model bicelle with the final curvature of 0.15 $nm^{-1}$ is shown. Initial planar bicelle is shown as pink lines. The final curved bicelle is light blue. Wall particles are shown as red (initial positions) and blue (final positions) spheres. The wall particles from lower wall, which are dropped after the bending, are gray. The arrows show direction of pulling of the wall particles. Green point indicates the center of curvature and the dashed lines indicate the sector occupied by the walls.

*Umbrella sampling protocol*

Several ligand molecules (five in case of water and ions and three in case of cisplatin and gemcitabine) were placed at equal intervals along the bilayer part of the bicelle. The harmonic biasing potential with the force constant of 1000 $kJ\cdot mol^{-1}\cdot nm^2$ was applied to the center of masses of each ligand along local Z axis (normal to the membrane plane at given point). The potentials were centered at discrete points distributed along local Z axis in 0.1 nm intervals. Additional weak flat-bottom potential was applied along X axis for each ligand to prevent their accidental interaction due to uncontrollable lateral diffusion. This produced 57 umbrella sampling windows spanning through the whole bilayer. The PMFs were computed as the average of all ligands using the weighted histogram technique as implemented in Gromacs [17].

Convergence of PMFs is monitored after each 3-4 ns of simulations. Simulations continue until systematic changes of the height and shape of the PMF occur between the check points. Equilibration time varies between the systems and ranges from 20 to 50 ns per window. A plot showing typical convergence of the free energy profile is shown in Figure S4.

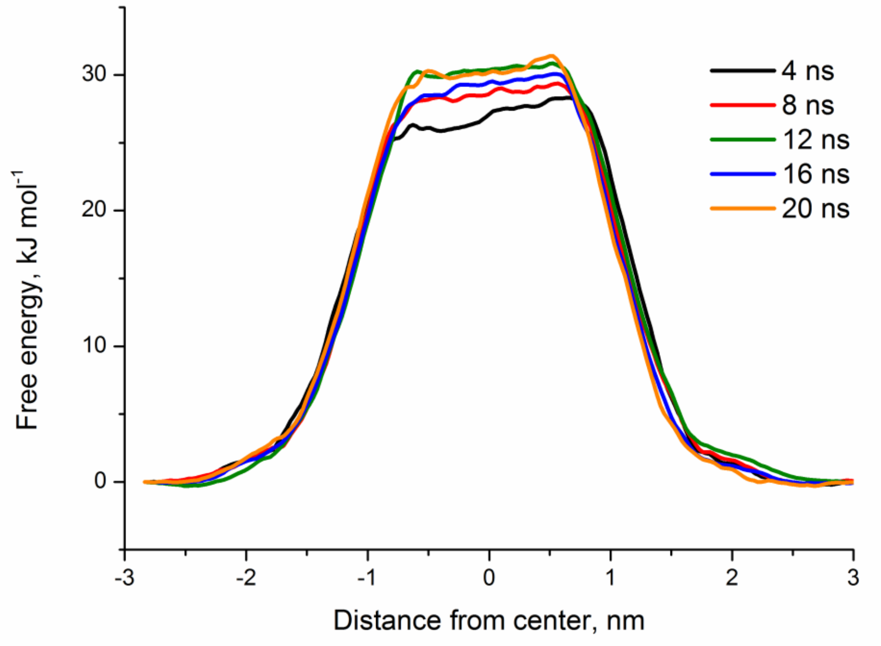


**Figure S4.** Example of the convergence of the PMF for water at zero curvature.

The errors of the PMFs were estimated using the bootstrapping method [18]. For each system 400 bootstraps were computed by considering complete histograms as independent data points using the Gromacs WHAM tool. Each PMF took ~1.5 μs of simulation time in average, which results in more than 20 μs of simulation time for all compounds and different values of curvature.

The diffusion coefficients D of the ligands were computed for each umbrella sampling window using the method of Hummer [19]:

$$D(z_i) = \frac{\langle \delta z^2 \rangle_i}{\tau_i},$$

where $\langle \delta z^2 \rangle_i$ and $\tau_i$ are mean square deviation from the average position and the position correlation time for the window *i*. The method of computing $\tau_i$ is based on Laplace transform of the position autocorrelation function. We used an implementation of this method proposed by DeMarco et. al. [20]. Diffusion coefficients of all ligands were averaged for each window. Obtained profiles of diffusion coefficient were further smoothed by the running average using the moving window of 3 points. The average of *D* for all the ligands from umbrella sampling windows *i*-1, *i* and *i*+1 is reported as the mean value of *D* for the window *i* and the standard deviation of these values is reported as the error of *D* for the window *i*.

The permeability coefficients *P* were computed using standard inhomogeneous solubility-diffusion model [21]:

$$P = 1 \Big/ \int_{z_1}^{z_2} R(z)dz,$$

where *R*(*z*) is the local permeation resistance of the membrane at depth *z* expressed as

$$R(z) = e^{\frac{W(z)}{kT}} \Big/ D(z)$$

where $W(z)$ is the PMF of a given ligand. The integration limits $z_1$ and $z_2$ were chosen as ±3 nm from the membrane center (the points in the water phase outside the membrane). The value of *P* is insensitive to the choice of these limits since the resistance *R* in water phase is negligible.

The errors of $R$ and $P$ could be evaluated using the error propagation formula based on the standard deviations of $W(z)$ and of $D(z)$ (namely $\sigma(W(z))$ and $\sigma(D(z))$). For the resistance $R$ at particular point $z$ we get:

$$\sigma(R(z)) = R(z)\sqrt{\left(\frac{\sigma(W(z))}{k_B T} e^{\frac{(\sigma(W(z))-W(z))}{k_B T}}\right)^2 + \left(\frac{\sigma(D(z))}{D(z)}\right)^2},$$

where $\sigma(R(z))$ is the standard deviation of R. In order to get global resistance one has to intergrate R(z) over z. Since the integral is approximated by a discrete sum the following expression for the standard deviation of R could be derived:

$$\sigma(R) = \sqrt{\sum_i (R_i dz_i)^2},$$

where i denotes one discrete value and dzi is the size of i-th integration step. Finally, the error of P is computed as:

$$\sigma(P) = P\frac{\sigma(R)}{R}.$$

*References*


1. Berendsen, H. J. C.; Marrink, S. J. Molecular dynamics of water transport through membranes: Water from solvent to solute. *Pure and Applied Chemistry* **1993,** *65* (12), 2513-2520.

2. Marrink, S.-J.; Berendsen, H. J. C. Simulation of water transport through a lipid membrane. *J. Phys. Chem.* **1994,** *98* (15), 4155-4168.

3. Lee, C. T.; Comer, J.; Herndon, C.; Leung, N.; Pavlova, A.; Swift, R. V.; Tung, C.; Rowley, C. N.; Amaro, R. E.; Chipot, C.; Wang, Y.; Gumbart, J. C. Simulation-Based Approaches for Determining Membrane Permeability of Small Compounds. *J. Chem. Inf. Model.* **2016,** *56* (4), 721-733.

4. Ribeiro, R. P.; Coimbra, J. T. S.; Ramos, M. J.; Fernandes, P. A. Diffusion of the small, very polar, drug piracetam through a lipid bilayer: an MD simulation study. *Theor Chem Acc* **2017,** *136* (4), 46.

5. Awoonor-Williams, E.; Rowley, C. N. Molecular simulation of nonfacilitated membrane permeation. *Biochimica et Biophysica Acta (BBA) - Biomembranes* **2016,** *1858* (7, Part B), 1672-1687.

6. Shinoda, W. Permeability across lipid membranes. *Biochim. Biophys. Acta* **2016,** *1858* (10), 2254-2265.

7. Dickson, C. J.; Hornak, V.; Pearlstein, R. A.; Duca, J. S. Structure–Kinetic Relationships of Passive Membrane Permeation from Multiscale Modeling. *J. Am. Chem. Soc.* **2017,** *139* (1), 442-452.

8. Swift, R. V.; Amaro, R. E. Back to the Future: Can Physical Models of Passive Membrane Permeability Help Reduce Drug Candidate Attrition and Move Us Beyond QSPR? *Chemical Biology & Drug Design 81* (1), 61-71.

9. Votapka, L. W.; Lee, C. T.; Amaro, R. E. Two Relations to Estimate Membrane Permeability Using Milestoning. *J. Phys. Chem. B* **2016,** *120* (33), 8606-8616.

10. Coimbra, J. T. S.; Fernandes, P. A.; Ramos, M. J. Revisiting Partition in Hydrated Bilayer Systems. *J. Chem. Theory Comput.* **2017,** *13* (5), 2290-2299.

11. Neale, C.; Bennett, W. F. D.; Tieleman, D. P.; Pomès, R. Statistical Convergence of Equilibrium Properties in Simulations of Molecular Solutes Embedded in Lipid Bilayers. *Journal of Chemical Theory and Computation* **2011,** *7* (12), 4175-4188.

12. Chipot, C.; Comer, J. Subdiffusion in Membrane Permeation of Small Molecules. *Scientific Reports* **2016,** *6*, 35913.

13. Gaalswyk, K.; Awoonor-Williams, E.; Rowley, C. N. Generalized Langevin Methods for Calculating Transmembrane Diffusivity. *Journal of Chemical Theory and Computation* **2016,** *12* (11), 5609-5619.

14. Yesylevskyy, S.; Ramseyer, C. Determination of mean and Gaussian curvatures of highly curved asymmetric lipid bilayers: the case study of the influence of cholesterol on the membrane shape *PCCP* **2014,** *16* (32), 17052-17061.

15. Yesylevskyy, S. O.; Kraszewski, S.; Ramseyer, C. Determination of the shape and curvature of nonplanar lipid bilayers that are bent in a single plane in molecular dynamics simulations. . *Journal of Molecular Modeling* **2014,** *20* (4), 2176.

16. Yesylevskyy, S. O.; Rivel, T.; Ramseyer, C. The influence of curvature on the properties of the plasma membrane. Insights from atomistic molecular dynamics simulations. *Scientific Reports* **2017,** *7* (1), 16078.

17. Hub, J.; de Groot, B.; van der Spoel, D. g_wham—A Free Weighted Histogram Analysis Implementation Including Robust Error and Autocorrelation Estimates. *Journal of Chemical Theory and Computation* **2010,** *6* (12), 3713-3720.

18. Hub, J.; De Groot, B.; der Spoel, D. g_wham-A free Weighted Histogram Analysis Implementation Including Robust Error and Autocorrelation Estimates. *J. Chem. Theory Comput.* **2010,** *6*, 3713-3720.

19. Hummer, G. Position-dependent diffusion coefficients and free energies from Bayesian analysis of equilibrium and replica molecular dynamics simulations. *New Journal of Physics* **2005,** *7*, 34-34.

20. DeMarco, K. R.; Bekker, S.; Clancy, C. E.; Noskov, S. Y.; Vorobyov, I. Digging into Lipid Membrane Permeation for Cardiac Ion Channel Blocker d-Sotalol with All-Atom Simulations. *Front. Pharmacol.* **2018,** *9* (26).

21. Marrink, S. J.; Berendsen, H. J. C. Permeation Process of Small Molecules across Lipid Membranes Studied by Molecular Dynamics Simulations. *J. Phys. Chem.* **1996,** *100* (41), 16729-16738.
